# The exchange dynamics of client molecules in biomolecular condensates

**DOI:** 10.64898/2026.07.06.736877

**Authors:** Ross Kliegman, Vladimir Grigorev, Yaojun Zhang

## Abstract

Biomolecular condensates are dynamic assemblies whose functions depend on continuous exchange of molecular components with the surrounding environment. While scaffold molecules drive phase separation and condensate architecture, many functional components are clients that are recruited through interactions with the scaffold-rich environment. Despite their prevalence, how client– scaffold interactions shape client exchange dynamics remains poorly understood. Here, we develop a reaction-diffusion model for client exchange in scaffold-driven condensates, in which clients switch between a scaffold-bound state and an unbound state. Bound clients exchange through scaffold-mediated transport, whereas unbound clients diffuse through the pore space of the condensate. Using the fluorescence recovery of fully photobleached condensates as a measure of client exchange, we compare transport through these two pathways with bound–unbound conversion and identify three limiting regimes. In the slow-conversion regime, bound and unbound clients recover through distinct scaffold- and pore-mediated pathways. In the intermediate-conversion regime, recovery of bound clients becomes limited by client unbinding. In the fast-conversion regime, local equilibrium between bound and unbound clients produces an effective single-state recovery. We further propose a unifying description that connects these regimes and quantitatively captures the apparent recovery timescales extracted from numerical simulations across condensate sizes. Our results provide a framework for interpreting component-specific exchange dynamics, and highlight client size, client– scaffold binding, and condensate porosity as key regulators of client turnover in multicomponent condensates.

## I. INTRODUCTION

Biomolecular condensates organize cellular biochemistry by concentrating selected proteins, nucleic acids, and metabolites into distinct compartments [1, 2]. Their functions depend not only on which molecules are enriched or excluded, but also on how rapidly these molecules interact with each other and are exchanged with the surrounding environment [3]. By tuning condensate dynamics, cells can regulate biochemical reaction rates, control the speed of condensate responses to signals, and modulate condensate number, size, and localization [4–7]. A quantitative understanding of the dynamical properties of condensates is therefore essential for elucidating their functions.

Condensate dynamics are routinely quantified by a variety of experimental techniques, such as fluorescence recovery after photobleaching (FRAP) [8, 9], single-molecule tracking [10, 11], fluorescence correlation spectroscopy [12, 13], and passive and active microrheology [14]. Complementary theoretical models have also been developed to interpret these measurements and extract physical quantities, such as diffusion coefficients, residence times, and exchange rates [8, 15–18]. While current models have provided important insights, they have largely focused on simplified condensates, often consisting of single molecular components. Cellular condensates, however, are generally composed of a multitude of components whose dynamics can differ substantially from one another [19, 20]. As quantitative measurements of multicomponent condensates become increasingly common, there is a growing need for theoretical models that can resolve and interpret the distinct dynamics of individual molecular components.

An important dynamical property of condensates is their exchange with the surrounding dilute phase, which controls molecular turnover and condensate responses to environmental changes. This exchange is commonly probed by full-FRAP experiments, in which fluorescence recovery after bleaching an entire condensate reports the timescale of molecular replacement by dilute-phase molecules [18, 21, 22]. In a multicomponent condensate, exchange can be component-specific, with different molecular species entering, moving within, and leaving the condensate through distinct pathways. Understanding such component-specific transport requires a framework that connects molecular identity to the corresponding physical pathways.

A useful starting point for modeling such component-specific exchange is the scaffold-client framework [23, 24]. Scaffold molecules contribute strongly to phase separation and condensate architecture, whereas client molecules are recruited through interactions with the scaffold-rich environment but do not substantially perturb the scaffold-driven phase equilibrium. This framework naturally motivates a component-specific description of condensate exchange dynamics. For example, multivalent interactions among scaffold molecules can give rise to a percolated network in the dense phase [25, 26], creating a heterogeneous environment with a characteristic mesh or pore size [27, 28]. In such an environment, different molecular components can bind to the network and diffuse with low mobility or unbind from the network and diffuse more rapidly through the mesh-work pores. How each component binds to the network, unbinds from it, and diffuses through the pore space can therefore give rise to distinct exchange pathways with the surrounding dilute phase.

Following this physical picture, recent work [29] introduced a reaction-diffusion model for scaffold exchange dynamics, in which scaffolds can switch between a low-mobility bound state and a high-mobility unbound state while exchanging with the surrounding dilute phase. Notably, the model predicted that transport through the pore space can accelerate scaffold exchange and that when exchange is limited by scaffold unbinding from the network, the exchange timescale becomes independent of condensate size. These predictions were supported by FRAP measurements on a synthetic DNA nanostar condensate system [29].

The results in Ref. [29] raise an important question: how broadly does pore-mediated exchange operate in biological condensates? Because biological condensates tyically have much higher molecular volume fractions than synthetic DNA nanostar condensates (10–30% [30] versus 1–5% [31, 32]), they are likely less porous. In such dense environments, the unbound scaffold population can be strongly suppressed because scaffolds are often much larger than the characteristic pore size [28]. Clients, however, provide a natural context in which pore-mediated exchange remains important. Clients are often smaller than scaffolds, allowing an appreciable unbound population to occupy the pore space even in condensates with little or no unbound scaffold population [28]. As shown in Fig. 1, clients can bind to scaffold molecules and move with scaffold dynamics—either with the scaffold network inside the condensate or with dilute-phase scaffolds outside—or dissociate and diffuse more rapidly through the condensate pore space and the surrounding dilute phase. In such a picture, the exchange dynamics of clients are governed not only by client-scaffold binding affinity, but also by scaffold mobility and porosity of the condensate.

**FIG. 1.**
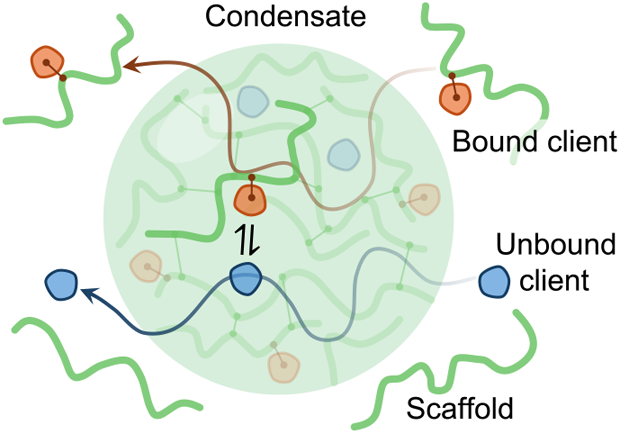
Schematic of client exchange in a multicomponent condensate. Clients can switch between scaffold-bound and unbound states, enabling slow scaffold-mediated exchange through bound-client motion and fast pore-mediated exchange through diffusion of unbound clients.

Here, we develop a reaction-diffusion model for client exchange in scaffold-driven condensates. We model clients as switching between scaffold-bound and unbound states, allowing client binding kinetics, scaffold mobility, and diffusion through pores to jointly determine client exchange, which we quantify via the full-FRAP recovery of photobleached condensates. By comparing the binding/unbinding timescale with the scaffold- and pore-mediated transport timescales, we identify distinct regimes in which client recovery is dominated by different rate-limiting processes and develop a unifying description that predicts the recovery timescales across regimes. We then use numerical reaction-diffusion simulations to validate these findings. Our work clarifies how client binding kinetics, scaffold mobility, and condensate porosity together shape component-specific turnover in multicomponent condensates.

## II. RESULTS

### A. Scaffold exchange dynamics

We first describe the exchange dynamics of scaffold molecules as a reference for the scaffold-associated contribution to client exchange. Following previous theoretical studies on scaffold exchange, we consider a spherical condensate of radius *R* with an equilibrium scaffold concentration profile *s*^eq^(*r*), where *r* is the distance from the condensate center. Assuming spherical symmetry, a minimal continuum description for the time evolution of the concentration of bleached scaffold molecules, *s*(*r, t*), is [15]:

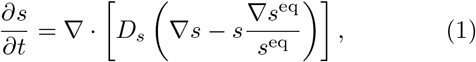

where *D*_*s*_(*r*) is the position-dependent scaffold diffusion coefficient. Eq. (1) describes the relaxation of the bleached scaffold population in a phase-separated environment, where the second term on the right-hand side represents the excess chemical potential that drives molecules toward a nonuniform equilibrium concentration profile.

For a full-FRAP experiment, the scaffold molecules inside the condensate are initially bleached while molecules outside are unbleached,

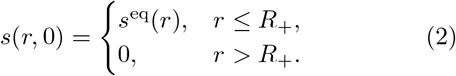

where *R*_+_ is the upper bound of the interface region. The concentration satisfies no-flux boundary conditions at the condensate center and system boundary: ∂*s*(*r, t*)*/*∂*r*|_*r*=0_ = ∂*s*(*r, t*)*/*∂*r*|_*r*→∞_ = 0. The corresponding normalized scaffold recovery curve is

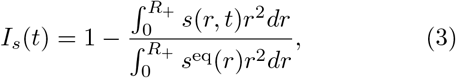

which reports the replacement of bleached scaffolds inside the condensate by unbleached scaffolds from the surrounding dilute phase.

The full-FRAP recovery time provides a means of measuring the exchange timescale associated with the dynamics in Eq. (1), which are mainly limited by three physical processes: diffusion in the dense phase, diffusion in the dilute phase, and transport across the interface. The interface contribution is normally much smaller than the other two, but can become rate-limiting when molecular transport across the interface is hindered by a local mobility minimum, a potential barrier, or frequent “bouncing” at the condensate interface [18, 33, 34]. Since interface resistance is not the focus of this work, we neglect interface contributions to both scaffold and client exchange for simplicity. The scaffold exchange timescale under the sharp-interface approximation is therefore [18]:

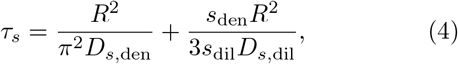

where *D*_*s*,den_ and *D*_*s*,dil_ are the scaffold diffusivities in the dense and dilute phases, respectively, and *s*_den_ and *s*_dil_ are the corresponding equilibrium concentrations.

### B. Client exchange dynamics

We next consider a client species that exchanges between the condensate and the surrounding dilute phase in the scaffold background. Assuming clients do not significantly perturb the scaffold equilibrium and dynamics, we model clients as switching between two states: a scaffold-bound state, with bleached concentration *c*_*b*_(*r, t*), and an unbound state, with bleached concentration *c*_*u*_(*r, t*). Bound clients move with scaffolds, whereas unbound clients diffuse freely.

Let 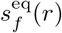 denote the equilibrium concentration of scaffolds available for client binding. We describe the dynamics of the bleached bound- and unbound-client populations using a reaction-diffusion model [29]:

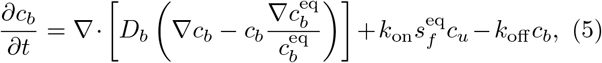

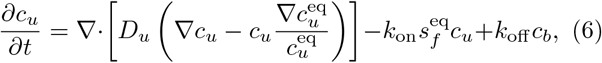

where *D*_*b*_(*r*) and *D*_*u*_(*r*) are the position-dependent diffusivities of bound and unbound clients, respectively, and 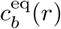 and 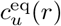 are their corresponding equilibrium concentration profiles. For bound clients, *D*_*b*_(*r*) = *D*_*s*_(*r*), reflecting their motion with scaffold dynamics. The first terms on the right-hand sides describe diffusion of the bound and unbound client populations. The second terms describe drift arising from the nonuniform thermodynamic environment, which drives each population toward its equilibrium spatial profile. The remaining terms describe local reversible binding and unbinding between clients and scaffolds,

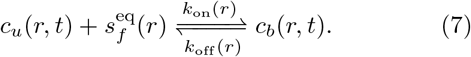

At equilibrium, local binding and unbinding must satisfy detailed balance, giving

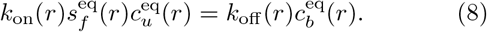

This condition allows the reaction terms to be written in terms of a single local conversion parameter,

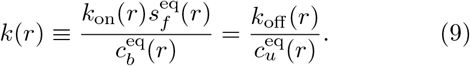

The reaction flux from the unbound to the bound state can then be written as 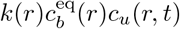, while the reverse flux is 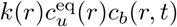. The client dynamics therefore become

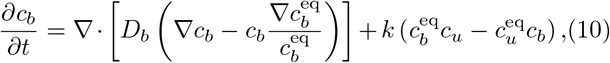

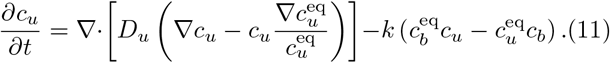

These equations make the equilibrium conditions explicit: the bound and unbound bleached populations relax to 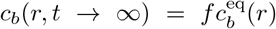 and 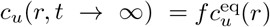, respectively, where *f* is the overall fraction of bleached clients in the system.

To model full FRAP of client molecules, we take both bound and unbound clients inside the condensate to be initially bleached,

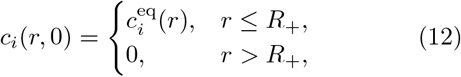

with *i* = *b, u*. Again, their concentrations satisfy no-flux boundary conditions at the condensate center and system boundary: ∂*c*_*i*_(*r, t*)*/*∂*r*| _*r*=0_ = ∂*c*_*i*_(*r, t*)*/*∂*r*| _*r*→∞_ = 0. The corresponding client recovery curve is

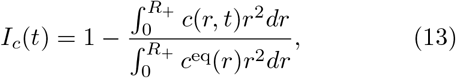

where

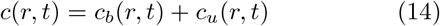

is the total concentration of bleached clients and

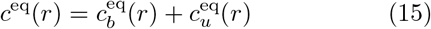

is the total equilibrium concentration of clients.

The coupled reaction-diffusion equations, Eqs. (10) and (11), reveal that client recovery is not controlled by a single transport process. Instead, the same client species can exchange through two coupled dynamical modes. The scaffold-bound population, *c*_*b*_, follows scaffold dynamics and therefore provides a scaffold-mediated pathway for recovery. The unbound population, *c*_*u*_, diffuses more rapidly through the condensate pore space and the surrounding dilute phase, providing a pore-mediated pathway. Because clients continuously convert between bound and unbound states, the overall recovery curve *I*_*c*_(*t*) reflects both transport of each species and conversion between them.

### C. Slow conversion reveals distinct scaffold- and pore-mediated exchange modes

The relative importance of transport and conversion depends on how the timescale of client-scaffold binding/unbinding compares with the transport timescales of the two pathways. We first explore the limit in which client-scaffold binding/unbinding is much slower than client exchange through both pathways. In this limit, conversion between bound and unbound clients becomes negligible over the recovery timescale, so the two populations contribute independently to the full-FRAP recovery. The scaffold-bound population recovers through a scaffold-mediated mode governed by scaffold dynamics, whereas the unbound population recovers through a pore-mediated mode governed by faster unbound-client diffusion.

The full-FRAP recovery curve can therefore be approximated as a sum of two components,

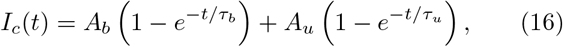

Where

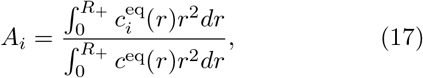

with *i* = *b, u. A*_*b*_ and *A*_*u*_ are the amplitudes of the scaffold-mediated and pore-mediated recovery modes, respectively, and *A*_*b*_ + *A*_*u*_ = 1. The corresponding recovery timescales can be derived in direct analogy with scaffold exchange

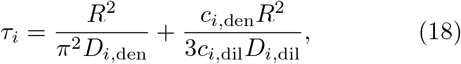

where *D*_*i*,den_ and *D*_*i*,dil_ are the diffusivities of state *i* in the dense and dilute phases, respectively, and *c*_*i*,den_ and *c*_*i*,dil_ are the corresponding equilibrium concentrations. Here, the scaffold-mediated recovery time, τ_*b*_, is controlled by the motion of scaffold-bound clients. Because bound clients remain stably associated with scaffolds over the recovery timescale, their equilibrium profile follows the scaffold equilibrium profile,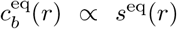, and their dynamics follows that of scaffolds, *D*_*b*_(*r*) = *D*_*s*_(*r*). The scaffold-mediated recovery time is therefore approximately equal to the scaffold recovery time, τ_*b*_ ≈τ_*s*_, in the slow-conversion limit.

In comparison, the pore-mediated recovery time, τ_*u*_, is controlled by unbound-client transport through the condensate pore space and the surrounding dilute phase. Because unbound clients are not constrained to move with the slowly rearranging scaffold network, their diffusion in the dense phase can be much faster than that of bound clients, *D*_*u*,den_ ≫*D*_*b*,den_. Unbound clients are also expected to partition much less strongly into the condensate than bound clients, *c*_*u*,den_*/c*_*u*,dil_ ≪*c*_*b*,den_*/c*_*b*,dil_. This follows because clients, by definition, partition much less strongly than scaffolds, (*c*_*b*,den_ + *c*_*u*,den_)*/*(*c*_*b*,dil_ + *c*_*u*,dil_) ≪*s*_den_*/s*_dil_, whereas bound clients follow the scaffold partitioning in this slow-conversion limit, *c*_*b*,den_*/c*_*b*,dil_ ≈*s*_den_*/s*_dil_. The pore-mediated recovery is therefore expected to be much faster than the scaffold-mediated recovery, τ_*u*_ ≪ τ_*b*_.

### D. Intermediate conversion shifts scaffold-mediated exchange to conversion-limited exchange

Because the pore-mediated recovery time is much shorter than the scaffold-mediated recovery time, τ_*u*_≪ τ_*b*_, there is an intermediate regime in which the client-scaffold conversion timescale lies between these two exchange timescales. In this regime, unbound clients can still exchange rapidly through the pore-mediated pathway, but scaffold-bound clients are not efficiently exchanged by scaffold-mediated transport. Instead, the bound population recovers by converting into the rapidly exchanging unbound population. As a result, the recovery curve *I*_*c*_(*t*) retains the two-component form, but the slow component is governed by a conversion-limited time rather than by the scaffold-mediated exchange time,

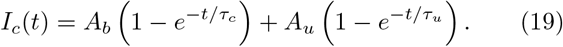

Because unbound clients exchange rapidly, the unbound population can be treated as quasi-equilibrated during the slower conversion process. The rate-limiting step for the slow mode is then the conversion of bound clients into the unbound state, followed by rapid replacement through the pore-mediated pathway. This gives a conversion-limited timescale [29]

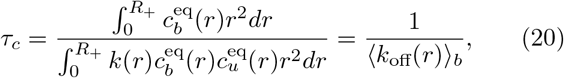

where⟨ · · ·⟩_*b*_ denotes an average over the equilibrium bound-client population within the condensate.

### E. Fast conversion yields an effective one-state description of client exchange

Finally, we explore the opposite limit in which the conversion is much faster than the exchange through either pathway. In this fast-conversion limit, the scaffold-bound and unbound client populations can be treated as being in local equilibrium during recovery,

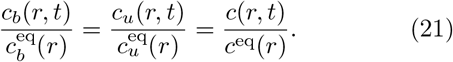

The local-equilibrium condition reduces the two coupled reaction-diffusion equations, Eqs. (10) and (11), to a single effective diffusion equation for the total bleached-client population,

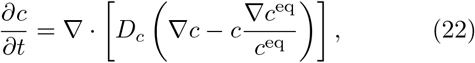

where *D*_*c*_(*r*) is an effective diffusion profile,

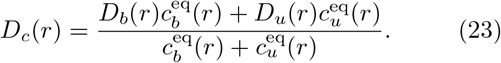

Thus, in the fast-conversion limit, client exchange is controlled by an equilibrium-weighted average of motion in the scaffold-bound and unbound states.

The effective single-field description in Eq. (22) leads to a single-mode of recovery with timescale,

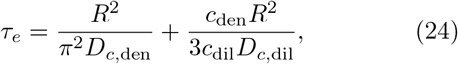

where *D*_*c*,den_ and *D*_*c*,dil_ are the effective client diffusivities in the dense and dilute phases, respectively, and *c*_den_ and *c*_dil_ are the corresponding total equilibrium client concentrations.

### F. A unifying description of client exchange across regimes

The analytical theory developed above identifies three limiting regimes depending on the ordering of the scaffold-mediated exchange time τ_*b*_, the pore-mediated exchange time τ_*u*_, and the conversion-limited time τ_*c*_.

(i) When conversion is slowest, τ_*u*_ ≪τ_*b*_ ≪ τ_*c*_, the two client populations recover independently, producing a two-component recovery curve in which the scaffold-mediated component relaxes with τ_*b*_ ≈τ_*s*_, whereas the pore-mediated component relaxes with τ_*u*_. (ii) In the intermediate regime, τ_*u*_ ≪ τ_*c*_ ≪ τ_*b*_, unbound clients still exchange rapidly through the pore-mediated pathway with τ_*u*_. However, bound clients recover primarily by converting into the rapidly exchanging unbound population, leading to a conversion-limited recovery with τ_*c*_. (iii) When conversion is fastest, τ_*c*_ ≪τ_*u*_ ≪ τ_*b*_, rapid interconversion drives bound and unbound clients into local equilibrium. The recovery curve is then approximately single-exponential, with the effective recovery time τ_*e*_.

To connect these regimes, we propose a unifying approximation for the full-FRAP recovery curve:

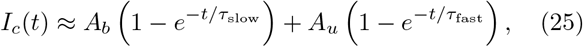

where τ_slow_ and τ_fast_ are the apparent slow and fast recovery times, respectively, obtained by interpolation between the limiting regimes,

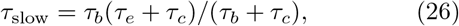

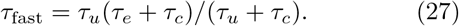

These expressions exhibit the expected limiting behaviors. In the slow-conversion regime, they reduce to τ_slow_ ≈τ_*b*_ and τ_fast_ ≈ τ_*u*_. In the intermediate regime, they reduce to τ_slow_ ≈τ_*c*_ and τ_fast_ ≈τ_*u*_. In the fast-conversion regime, both branches approach the effective recovery time, τ_slow_ ≈τ_fast_ ≈τ_*e*_, so the recovery curve becomes approximately single-exponential.

### G. Numerical simulations

To test these predictions for biological condensates, we specified radial profiles for the equilibrium concentrations and diffusivities of both bound and unbound client populations with a smooth transition across the interface [35]:

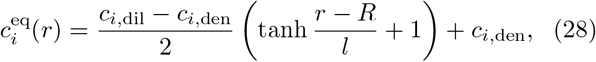

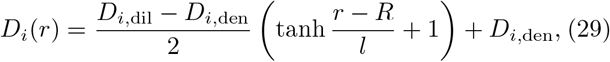

where *i* = *b, u* and *l* sets the interface width (Figs. 2a and 2b). We note that these profile choices naturally make the interface contributions to the exchange timescales negligible [18]. For the binding-unbinding kinetics, we assumed a spatially uniform unbinding rate, *k*_off_, which is physically motivated since unbinding primarily reflects the intrinsic lifetime of a bound client and is expected to be less sensitive to the surrounding environment. The local conversion coefficient in Eqs. (10) and (11) is then dtermined by detailed balance as 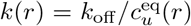. The conversion timescale becomes τ_*c*_ = 1*/k*_off_ with this choice of *k*(*r*). After these specifications, we chose parameters (Table I) such that the relevant timescales are well separated, allowing the slow-, intermediate-, and fast-conversion limits to be tested separately.

**FIG. 2.**
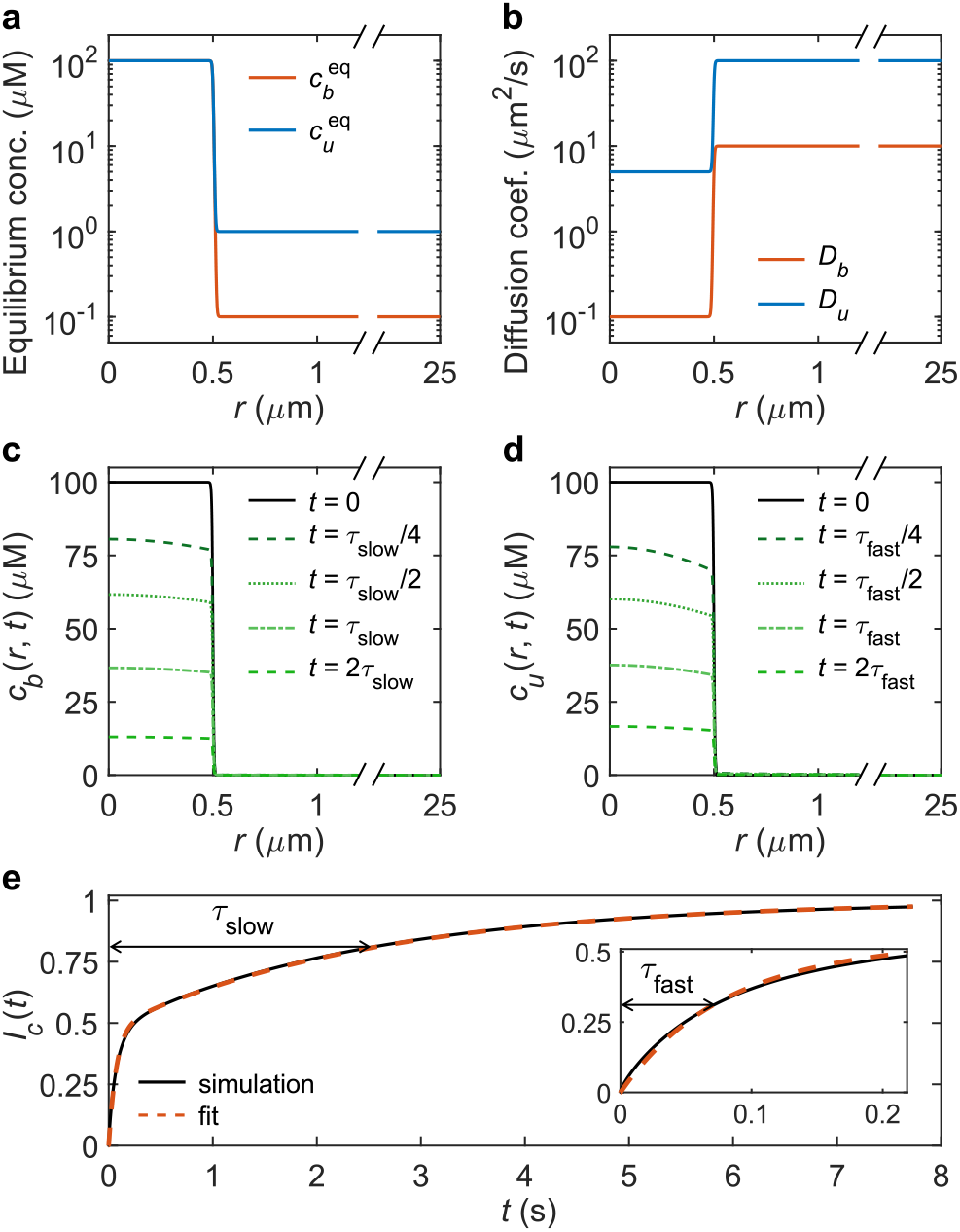
Representative numerical simulation of client exchange dynamics for a condensate of radius *R* = 0.5 *µ*m (parameters from Table I). (a,b) Equilibrium concentration profiles and diffusion-coefficient profiles of bound and unbound clients. (c,d) Time evolution of the initially bleached bound- and unbound-client concentration profiles after full-condensate photobleaching. The bound- and unbound-client profiles are shown at times scaled by the fitted slow and fast recovery timescales, τ_slow_ and τ_fast_, respectively. (e) Normalized recovery curve *I*_*c*_(*t*) from the numerical simulation and fit to Eq. (25), with the inset highlighting the early-time recovery. The extracted parameters, *A*_*b*_ = 0.52, *A*_*u*_ = 0.48, τ_slow_ = 2.52 s, and τ_fast_ = 0.073 s, agree well with their corresponding theoretical expectations, *A*_*b*_ = *A*_*u*_ = 0.5, τ_slow_ = 2.53 s, and τ_fast_ = 0.091 s.

**TABLE 1.** Parameters for numerical simulations

| $c_{b,\text{den}}$ | $c_{b,\text{dil}}$ | $c_{u,\text{den}}$ | $c_{u,\text{dil}}$ | $l$ |
| --- | --- | --- | --- | --- |
| 100 $\mu\text{M}$ | 0.1 $\mu\text{M}$ | 100 $\mu\text{M}$ | 1 $\mu\text{M}$ | 0.01 $R$ |
| $D_{b,\text{den}}$ | $D_{b,\text{dil}}$ | $D_{u,\text{den}}$ | $D_{u,\text{dil}}$ | $k_{\text{off}}$ |
| 0.1 $\mu\text{m}^2\text{s}^{-1}$ | 10 $\mu\text{m}^2\text{s}^{-1}$ | 5 $\mu\text{m}^2\text{s}^{-1}$ | 100 $\mu\text{m}^2\text{s}^{-1}$ | 0.3 $\text{s}^{-1}$ |

We implemented the numerical simulations by solving the two-state reaction-diffusion framework, Eqs. (10) and (11), under spherical symmetry using the full-FRAP initial condition in Eq. (12) and no-flux boundary conditions at the condensate center and the outer boundary of the simulation domain. The outer boundary was placed sufficiently far so that there were enough unbleached dilute-phase molecules for complete FRAP recovery. After obtaining the numerical solutions for *c*_*b*_(*r, t*) and *c*_*u*_(*r, t*) (Figs. 2c and 2d), we computed the full-FRAP recovery curve *I*_*c*_(*t*) using Eq. (13) with *R*_+_ = *R*+2*l* (Fig. 2e). The resulting recovery curves were fitted to Eq. (25) to extract the relevant amplitudes and timescales, which were then compared with the analytical predictions. See Methods for details of the numerical simulations and fitting procedures.

Because τ_*b*_ and τ_*u*_ scale as *R*^2^, whereas τ_*c*_ = 1*/k*_off_ is independent of *R*, varying the condensate size naturally allows the system to pass through the three limiting regimes (Fig. 3). For a small condensate with *R* = 0.125 *µ*m, the timescales satisfy τ_*u*_ ≪τ_*b*_ ≪τ_*c*_, placing the system in the slow-conversion regime (i). The recovery is therefore characterized by two apparent timescales: the fitted slow timescale τ_slow_ is close to the scaffold-mediated time τ_*b*_, whereas the fitted fast timescale τ_fast_ is close to the pore-mediated time τ_*u*_ (Fig. 3). For an intermediate-sized condensate with *R* = 1 *µ*m, the ordering becomes τ_*u*_ ≪τ_*c*_ ≪τ_*b*_, corresponding to the intermediate-conversion regime (ii). The recovery is still characterized by two apparent timescales: the fitted τ_fast_ remains close to the pore-mediated time τ_*u*_, but the fitted τ_slow_ is now close to the conversion-limited time τ_*c*_ (Fig. 3). For a large condensate with *R* = 8 *µ*m, the ordering becomes τ_*c*_ ≪τ_*u*_≪ τ_*b*_, corresponding to the fast-conversion regime (iii). In this regime, the fitted τ_slow_ and τ_fast_ collapse toward the same value, both converging to the effective recovery time τ_*e*_ (Fig. 3). Overall, across the full range of condensate sizes, the fitted apparent timescales follow the crossover behavior predicted by Eqs. (26) and (27) (Fig. 3).

**FIG. 3.**
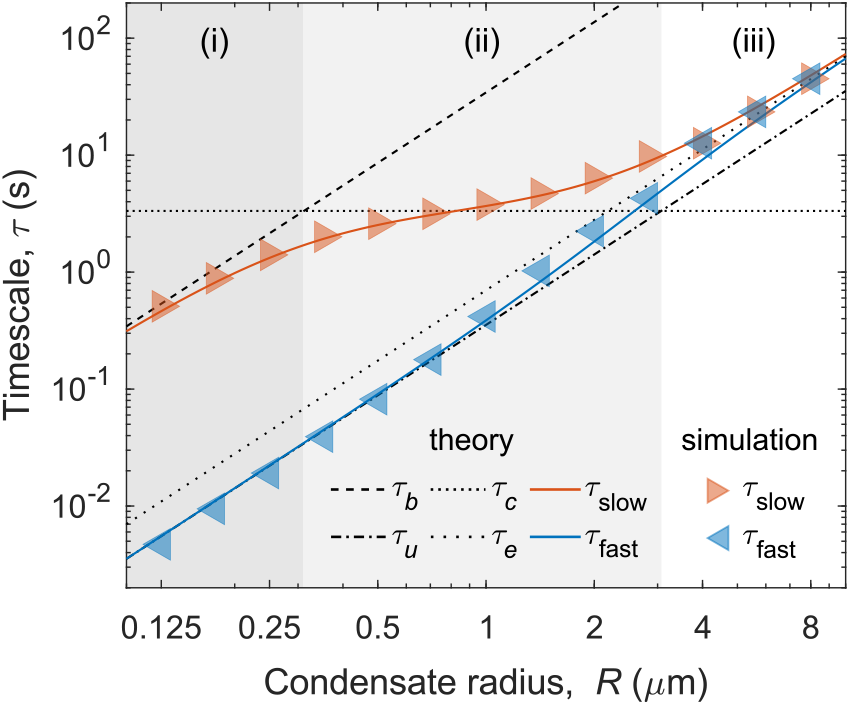
Numerical demonstration of client exchange timescales across regimes (parameters from Table I). Black dashed and dotted lines show the characteristic timescales for scaffold-mediated exchange, τ_*b*_, pore-mediated exchange, τ_*u*_, bound–unbound client conversion, τ_*c*_, and effective single-state exchange, τ_*e*_. As the condensate radius increases, the system transitions from (i) slow conversion, τ_*u*_ ≪τ_*b*_ *<* τ_*c*_ (dark gray), to (ii) intermediate conversion, τ_*u*_ *<* τ_*c*_ *<* τ_*b*_ (light gray), and finally to (iii) fast conversion, τ_*c*_ *<* τ_*u*_ ≪τ_*b*_ (unshaded). Colored solid curves show the predicted apparent recovery times, τ_slow_ and τ_fast_, from Eqs. (26) and (27), and symbols show the corresponding timescales extracted by fitting numerical full-FRAP recovery curves.

## III. DISCUSSION

We have developed theory and simulations to understand the exchange dynamics of client molecules in scaffold-driven biomolecular condensates. The central idea is that clients can exchange with the surrounding dilute phase through two coupled pathways. In the scaffold-mediated pathway, clients bind to scaffold molecules and inherit their slower dynamics, whereas in the pore-mediated pathway, clients dissociate from the scaffold and diffuse more rapidly through the condensate pore space and the surrounding dilute phase. These two pathways are coupled by reversible client–scaffold binding and unbinding, so transport through one state can feed recovery of the other. This physical picture naturally leads to component-specific exchange dynamics: clients can exchange on timescales that differ substantially from those of the scaffold.

What determines the balance between the scaffold- and pore-mediated transport? One key factor is the equilibrium partitioning of clients between the scaffold-bound and unbound states, which is controlled by client– scaffold binding affinity, the availability of binding sites, and the pore space accessible to unbound clients. This partitioning sets the relative amplitudes of the two pathways: stronger binding or more accessible binding sites favors the scaffold-bound population, whereas greater pore-space accessibility favors the unbound population. The balance between these transport pathways also depends on how much faster unbound clients diffuse than bound clients and how rapidly clients switch between the two states. Slow scaffold dynamics impede the mobility of bound clients and thus suppress direct scaffold-mediated exchange, whereas high condensate porosity and small client size facilitate diffusion of unbound clients through the pore space, promoting pore-mediated transport. The bound–unbound conversion rate then determines how strongly transport through the two pathways is coupled.

The physical picture proposed here may have implications for condensate function. Small molecules such as metabolites, short peptides, and small folded domains can enter condensates through the pore space and exchange rapidly, whereas larger molecules such as multivalent protein complexes, long RNAs, and ribonucleoprotein assemblies are more likely to be sterically hindered from accessing the pore space [28], whose exchange therefore depends more on interfacial capture, scaffold remodeling, or active cellular processes. Thus, the same condensate can behave as a rapidly accessible medium for small molecules while acting as a slow-retention compartment for larger molecules. Such size-dependent exchange could allow condensates to retain enzymes or regulatory factors while permitting substrates, products, and small cofactors to turn over efficiently.

Our model is intentionally minimal to keep the physical mechanisms transparent and the mathematical structure analytically tractable. Several extensions are natural. First, clients may engage in multivalent interactions with multiple scaffold molecules, making their motion decoupled from that of a single scaffold molecule and leading to a spectrum of binding and mobility states rather than switching between two discrete states. Second, scaffold networks may be spatially heterogeneous, causing binding-site density, local mobility, and pore accessibility to vary across the condensate [36, 37]. Such heterogeneity could create spatially dependent exchange pathways, with some regions favoring scaffold-associated retention and others favoring rapid pore-mediated transport. Finally, cellular condensates are often maintained away from equilibrium by active processes such as enzymatic remodeling, ATP-dependent reactions, RNA processing, or molecular synthesis and degradation [19, 38, 39]. Incorporating these effects will be important for connecting the minimal two-pathway picture developed here to the diverse exchange behaviors observed in living cells.

### IV. METHODS

### A. Numerical details for solving *c*_*b*_(*r, t*), *c*_*u*_(*r, t*), **and** *I*_*c*_(*t*)

For each choice of condensate radius *R*, we numerically solved the coupled reaction-diffusion equations, Eqs. (10) and (11), using MATLAB’s *pdepe* function [40, 41]. The *pdepe* solver computes solutions to initial-boundary-value problems for systems of parabolic partial differential equations (PDEs) in one spatial variable and time, with the symmetry parameter *m* = 2 for spherical geometry. The solver discretizes the spatial coordinate on a specified mesh and integrates the resulting system of ordinary differential equations using a variable-step, variable-order time integrator.

Specifically, we used a piecewise-defined radial mesh: a coarse spacing Δ*r*_bulk_ = *l/*2 inside and outside the condensate (0 ≤*r* ≤*R*_−_ and *R*_+_ ≤ *r* ≤ *R*_box_), and a fine spacing Δ*r*_int_ = *l/*5 in the interfacial region (*R*_−_ *< r < R*_+_), with *R*_±_ = *R* ± 2*l*. Here, *R*_box_ denotes the outer boundary of the simulation domain and was set to *R*_box_ = 50*R* for all simulations. Simulations were performed over the time interval 0 ≤*t* ≤*T*, where *T* = 3τ_slow_ and τ_slow_ was predicted by Eq. (26) for each choice of *R*. The numerical solutions for the bleached bound- and unbound-client concentrations, *c*_*b*_(*r, t*) and *c*_*u*_(*r, t*), were recorded at 1000 time points evenly spaced on a logarithmic scale between 10^−5^ s and *T* .

After obtaining *c*_*b*_(*r, t*) and *c*_*u*_(*r, t*), we computed the normalized full-FRAP recovery curve *I*_*c*_(*t*) according to Eq. (13). The radial integrals were evaluated numerically on the same grid used for the PDE solution.

### B. Fitting procedures for extracting τ_slow_ and τ_fast_

To extract the recovery amplitudes and timescales from the simulated full-FRAP curves, we fitted the numerical *I*_*c*_(*t*) to

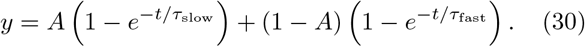

To reduce the influence of higher-order modes at early times, fitting was performed on log [1 − *I*_*c*_(*t*)] rather than directly on *I*_*c*_(*t*). For small condensates, where the two relaxation modes are well separated in time, all three fitting parameters, *A*, τ_slow_, and τ_fast_, can be reliably extracted, and the fitted amplitudes were close to the theoretical expectation *A* = 0.5. For example, the representative simulation shown in Fig. 2 gave *A* = 0.52. However, for large condensates, where τ_slow_ and τ_fast_ approach each other, the fitted amplitude *A* can show large variations due to parameter degeneracy. Therefore, for the results shown in Fig. 3, we fixed *A* = 0.5 and fitted only τ_slow_ and τ_fast_. In the fast-conversion regime, we further constrained τ_slow_ and τ_fast_ to have the same value, τ_eff_, so that 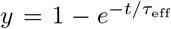 . The theoretical estimates τ_*b*_, τ_*u*_, and τ_*e*_ were used as initial guesses for τ_slow_, τ_fast_, and τ_eff_, respectively.

## DATA AVAILABILITY

The data underlying this article were generated from the simulations described in the text. The MATLAB scripts used to generate, analyze, and plot these data are available at [repository link].

## ACKNOWLEDGMENTS

We are grateful to … for insightful discussions and valuable feedback on the manuscript. This work was supported by a startup fund at Johns Hopkins University, NIH Award R35GM162296, and the Alfred P. Sloan Foundation through a Sloan Research Fellowship to Y.Z. (FG-2025-25076).

